# Environmental and demographic mechanisms provide relative stability for a vulnerable island songbird in novel conditions

**DOI:** 10.1101/2025.03.11.642629

**Authors:** James C. Mouton, Mario Pesendorfer, T. Scott Sillett

## Abstract

Understanding demographic and ecological mechanisms underlying population dynamics is a key goal in population ecology and critical for effective management in the face of climate change. Species may be well adapted to persist under normal ranges of environmental conditions, but increasingly novel environmental conditions may surpass the demographic buffering mechanisms in many populations. Small, isolated populations, such as those on islands, are expected to be especially vulnerable to declines caused by novel environmental conditions. We used an integrated population model to (1) examine ecological drivers of population growth and (2) assess global population trends of the Island Scrub-Jay (*Aphelocoma insularis*) from 2009-2019. Our results suggest that population size increased slightly over this interval despite declines during severe drought. We also found evidence that density dependence, precipitation, and food availability affected fecundity and the survival of non-breeding individuals. Breeder survival was relatively stable and had a weak effect on population growth as expected for long-lived species. Overall, our results provide an optimistic snapshot for this species by demonstrating resilience to contemporary drought, but also emphasize the species’ potential vulnerability due to its small population size.

## INTRODUCTION

Theory predicts that life history traits most strongly linked to population growth should be adapted to buffer population growth from environmental variation typically experienced by a species (Pfister 1998; Hilde et al. 2020; Le Coeur et al. 2022). For example, species with strong density dependence and slower life histories have evolved strategies to limit environmentally induced variation in adult survival, a trait with an outsized impact on fitness and population growth (mammals: Gaillard et al. 1998; reptiles: Miller et al. 2011; birds: Sæther et al. 2002; Martin & Mouton 2020). Recent climate change and human activity has forced natural populations to confront increasingly novel environmental conditions caused by extreme weather, invasive species, and habitat degradation (Sih et al. 2011). Such novel conditions may exceed a population’s capacity to buffer important vital rates that evolved under more typical environments, leading to declines in abundance (Schlaepfer et al. 2002; Ghalambor et al. 2007). Small, isolated populations, such as those on islands, are expected to be especially vulnerable to declines caused by novel environmental conditions (Quinn & Hastings 1987; Otto et al. 2017). Thus, studies examining how different forms of environmental variation affect vital rates of insular populations are critical for understanding population-level responses and devising effective management strategies.

Island species generally exhibit slower life history strategies, such as greater survival rates, delayed breeding, reduced fecundity, and strong density dependence due to limited habitat availability (MacArthur & Wilson 1967; McCallum et al. 2000; Covas 2012; Beauchamps 2021; Jezierski et al. 2023). These traits may allow island populations to be robust to environmental variation in multiple ways. First, delayed breeding in saturated habitats can create a persistent, non-territorial sub-population. These reproductively mature “floaters” can transition to breeders when territories become available, buffering losses of breeding individuals and enabling more consistent reproductive potential at the population level (Penteriani et al. 2011; Robles & Ciudad 2020). Second, strong density dependence facilitates rapid recovery of island populations following catastrophes such as storms, drought, or disease because vital rates increase at low when population density (McCallum et al. 2000; Seavy et al. 2009). Finally, slower life histories may favor behavioral plasticity that allows individuals to allocate resources towards survival during lean years and to focus on reproduction in years with abundant resources (Gibbs & Grant 1987; Maspoons et al. 2019; Martin & Mouton 2020). Island species with slow life history strategies are expected to have population dynamics strongly driven by variation in fecundity and non-breeder survival but relatively stable breeder survival (Gaillard et al. 1998; Hilde et al. 2020). Taken together, delayed breeding, density dependence, and slower life histories are expected to buffer population growth and increase the likelihood of persistence against environmental variation.

Extreme environmental conditions can strain populations’ capacity to buffer demographic change. For example, heat waves due to climate change are increasingly surpassing physiological limits, resulting in high adult mortality and declines across taxa (Ruthrof et al. 2018; Ratnayake et al. 2019; Riddell et al. 2019). Climate change is similarly increasing the frequency and duration of droughts that reduce survival and opportunities for compensatory reproduction (Prugh et al. 2018). Extreme weather may also interact with other novel conditions, such as invasive disease outbreaks, to erode fecundity and breeder survival with severe implications for population dynamics (Bakker et al. 2020, Rodríguez-Caro et al. 2021). Anamalous weather can also stress key biotic interactions with critical implications for population dynamics (Prugh et al. 2018). For example, the availability of important food sources can vary with climate and may have either immediate or lagged effects on populations of consumers (Boag & Grant 1981; Prugh et al. 2018). Extreme variation in predator densities driven by natural or human-caused factors may also limit the efficacy of demographic buffering among prey and lead to instability. Ultimately, determining the environmental and demographic mechanisms underlying contemporary population dynamics is essential for understanding the capacity of populations to buffer environmental variation and predicting population-level responses to future climate change.

Here we use the Island Scrub-Jay (*Aphelocoma insularis;* hereafter jays), a species of conservation concern, as a model species to examine these ideas. This species is endemic to 250 km^2^ Santa Cruz Island off the coast of Southern California, USA. The IUCN Red List uses an obsolete taxonomy the classifies all California *Aphelocoma* as *A. californica* (California scrub-jay), a species of “Least Concern.” However, *A. insularis* is listed as “Vulnerable” in an earlier Red List version (e.g., Sillett et al. 2012), and multiple studies have documented the species’ strong genetic and phenotypic differentiation and small population size (e.g., Delany and Wayne 2005, Sillett et al. 2012, Cheek et al. 2022, DeRaad et al. 2022). Over-grazing by sheep, deer, and elk, as well as rooting by feral pigs severely degraded the chaparral and woodland habitat used by jays on the island over the last two centuries, potentially resulting in population declines (Morrison et al. 2011; Sillett et al. 2012; Mosher et al. 2021). These invasive herbivores were eradicated by 2007, allowing scrub and understory vegetation to begin to return across much of the island (Beltran et al. 2014; Morrison et al. 2016) improving habitat conditions for jays. A recent study suggests that survival of breeding jays and population size may have plummeted from 2000-2006 (Mosher et al. 2021). However, whether these declines occurred at island-wide spatial scales and represent an ongoing threat to the species in the long-term are unclear.

We use an integrated population model to assess the demographic and environmental drivers of jay population dynamics from 2009-2019. We examined the effects of abiotic (precipitation, temperature) and biotic (population density, food availability, predator abundance) conditions on jay population dynamics. Our model integrates data from an island-wide point count survey (2009; 307 sites), detailed capture-recapture and fecundity data (2009-2019; 6 plots across the island), and a separate island-wide point count surveys of terrestrial birds (2013-2019; 120 sites). Our goals were (1) to assess the importance of density dependence in regulating the population, (2) to test *a priori* hypotheses about how key environmental factors like food availability, climate, and predator abundane drive variation in vital rates, and (3) to assess trends in vital rates and the island-wide population size of jays. Specifically, we tested whether abiotic factors like precipitation and temperature extremes are important drivers of jay population dynamics. Precipitation in fall and winter can have major effects on primary productivity and food availability for jays which could shape patterns in survival of breeders and floaters as well as fecundity. We tested for short-term and lagged effects of precipitation since rainfall has caused lagged effects in some species on the adjacent mainland (Sakai 2016; Prugh et al. 2018). Extreme temperatures can also impact survival by increasing metabolic costs or directly causing mortality (Swanson & Olmstead 1999; McKechnie & Wolf 2010). Thus, we tested whether maximum summer temperatures or minimum winter temperatures affected breeder or floater survival. We also considered key variation in biotic variables including food availability and predator density. We tested whether the abundance of acorns, a main food source during fall, improved annual survival of breeders or non-territorial floaters. Acorn abundance could also improve reproductive output during the following breeding seasons through improved body condition of jays prior to breeding or by greater food availability during the breeding season dues to more acorns being stored in caches (Koenig et al. 2009). These ideas assume that acorn abundance has carry-over effects on jay vital rates, but breeding jays may also be able to detect cues that predict the acorn crop the following fall. Indeed, other seed-eating species are known to anticipate seed resource pulses and increase reproductive output to match future seed abundance (Boutin et al. 2006).Thus, jays might also adjust reproductive effort so that they produced more offspring in years when they will experience greater acorn abundance and thus survival in their first fall. Finally, nest predation can be a major driver of fecundity, so we tested whether the density of island fox (*Urocyon littoralis*), a known predator of jay nests, influenced patterns in fecundity across years (Caldwell et al. 2012).

## METHODS

### Study Area

Santa Cruz Island is a 250 km^2^ island 32 km off the coast of southern California, USA (34.0022°N, -119.7225°W). It has a mediterranean climate with most of the rainfall occurring in winter followed by warm, dry summers. Habitat types on the island consists of approximately 54% scrub (e.g. coastal sage scrub), 23% oak chaparral, 6% forest, 10% grass and forb dominated habitat types. The terrain on the island is rugged and mountainous with elevation spanning from sea level to 740 m.

### Data collection

#### Population Surveys

Distinct point count surveys were conducted in 2009 and from 2013-2019. In April 2009, point count surveys were conducted at 307 locations spread across Santa Cruz Island (Fig. 1; Sillett et al. 2012). This study yielded an island-wide estimate of jay population size (1705; 95% CI = 1212-2369) which we incorporated directly into our model (see below). From 2013-2019, annual spring land bird surveys were conducted using protocols created by Channel Islands National Park. In each year, 120 survey locations were sampled mostly along the east-west axis of the island (Fig 1). Field technicians hiked to each survey point from the road and conducted unlimited radius point count surveys for 10 mins. The species, distance from the observer and the time-of-detection for each individual bird was recorded. We only considered detections within 300m from the observer (Sillett et al. 2012).

**Fig. 1:**
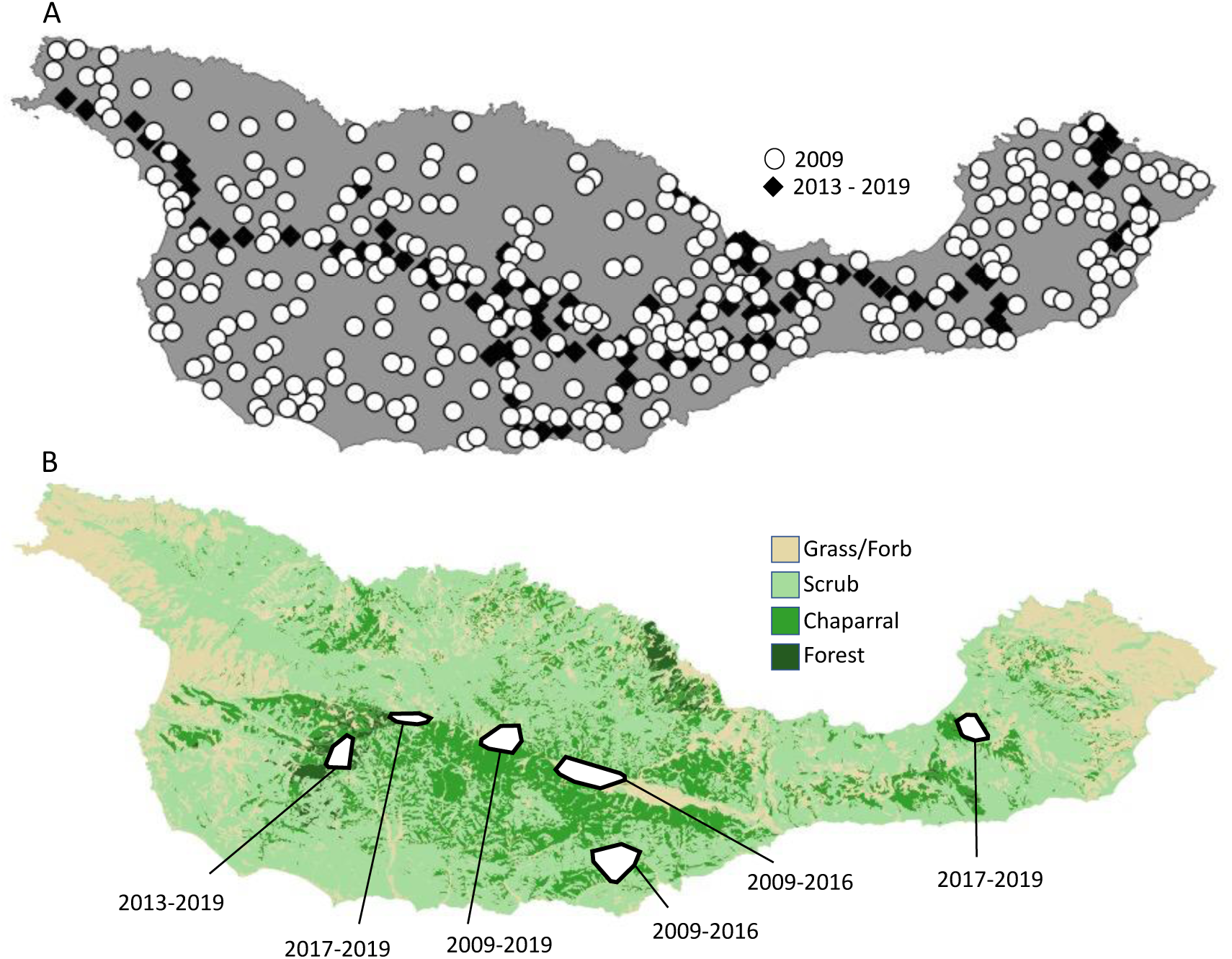
Maps of Santa Cruz Island showing **A)** survey sites from 2009 (white circles) and 2013-2019 (black diamonds), and **B)** the location of 6 jay demographic study plots and the time range we were actively collecting capture-recapture and/or reproductive data at each.

#### Capture-Recapture Data

We captured, marked and resighted jays during fall and spring from 2009-2019. We captured jays on each plot primarily using box traps baited with peanuts. We marked birds with numbered aluminum and unique combinations of plastic color leg bands (Caldwell et al. 2013). We monitored and resighted breeding pairs and floaters during nest searching and territory mapping activities. Resighting and capture efforts occurred primarily from Oct-Nov in the fall season and February-June during the spring season.

#### Fecundity Data

From 2009-2019, we conducted detailed demographic studies of breeding jays from approximately February to June. We followed territorial pairs, mapping the extent of territories, finding nests, and counting the number of young successfully fledged from each nest. Because renesting after nest failure is common, most pairs had multiple nesting attempts per season. In general, jays will only raise one successful brood per season, with rare exception (Delaney & Cheek 2022). We used the total number of fledglings for those territories for which we are confident we observed all nesting attempts and young.

#### Environmental Data

Environmental data came from several different sources. Precipitation and temperature data were collected on Santa Cruz Island and provided by the Western Regional Climate Center (WRCC 2023). We calculated the total rainfall during the “bioyear” between July 1 and June 30 to capture the precipitation in the year leading up to each breeding season. Island fox population size estimates were provided by the long-term monitoring program of The Nature Conservancy (Bakker et al. 2005). We investigated the acorn production of Santa Cruz Island oaks on a transect across a majority of the east-west extent of the island (*Quercus pacifica* [n=138] and *Q. agrifolia* [n=53]; Pesendorfer et al. 2014, 2018) during autumn from 2008-2019. To determine annual acorn production of individually-marked trees, we used the canopy survey method (Koenig et al. 1994). Two observers scanned the canopy of the tree for 15 seconds and counted all visible acorns. Each observers counts were combined and we used the mean acorn count across all trees as our index of annual acorn abundance.

### Integrated Population Model

We used an integrated population model (Zipkin & Saunders 2018; Schaub & Kéry 2022; hereafter IPM) to jointly analyze data on abundance from surveys at the population-level together with data on survival and reproduction from the detailed study of marked individuals. Analyzing submodels for these datasets together with a population model in an IPM allowed us to generate more precise estimates of demographic parameters, including population trends due to distinct information about vital rates provided by each dataset (Zipkin & Saunders 2018; Zipkin et al. 2019; Schaub & Kéry 2022). We used a Bayesian framework which allowed us to propagate uncertainty and facilitated the inclusion of incomplete data or datasets collected under distinct protocols (Schaub & Kéry 2022)

#### Count sub-model

We modelled count data from 2013-2019 using a normal distribution and assumed that the annual point counts surveys sampled a constant proportion of the island-wide population across years. We used an informative beta prior distribution (α = 2, β = 20) centered on ratio of the average number of birds counted across the surveyed sites and the island-wide population size from 2009 (Sillett et al. 2012) to model this proportion. We then used the product of this proportion and the island-wide population size to model the portion of the population available to be sampled during point count surveys. We used a weakly informative gamma distribution (α = 1.5, β =0.01) to model error in this sub-model.

#### Fecundity sub-model

We modelled the number of fledglings from 2009-2019 using a zero-inflated Poisson distribution. We included random effects for each plot and year for both the mean and proportion of zero parameters. We used informative priors for the average number of fledglings (normal: mean = 0.9, sd = 0.5) and proportion of zero (beta: α = 7, β = 3) parameters based on previous work. We modelled the average number of fledglings produced island-wide each year as the product of the average number of fledglings per female with the total number of breeders. The actual number of fledglings produced each year was modelled using a Poisson distribution to incorporate demographic stochasticity.

#### Survival sub-model

We analyzed capture-recapture data from 2009-2019 with a Cormack-Jolly-Seber model. We modelled survival across years and seasons (fall-to-spring and spring-to-fall) and included fixed terms for breeding status (breeder vs. floater). We included fixed effect terms for breeding status, transient effects, and sex, and allowed detection probability to vary across years and seasons. Because some plots stopped being sampled in some years, we also accounted for variation in sampling effort among plots using a fixed factor indicating whether the plot a bird was initially captured on was sampled in any given year. We calculated annual survival probabilities for floater and breeders to use in the population model. We used informative prior distributions for survival on the logit scale (median = 0.78; mode = 0.94; normal: 1.25, sd = 1.3) based on previous work (Bakker et al. 2020).

#### Population model

We integrated abundance, fecundity, and survival sub-models to model island-wide population size from 2009-2019. We parameterized island-wide population size in 2009 using a negative binomial prior distribution matching estimates from previous work (1705; 1212-2369 95%CI; Sillett et al. 2012). We modelled the number of surviving juvenile (from fledge until the subsequent breeding season), floater, and breeder survival using binomial distributions and survival parameters. We assumed similarly high survival among juveniles as older floaters based on previous work (Bakker et al. 2020). Breeder birds typically maintain the same territories for their whole lives and breeding territories are thought to be limited (Delaney & Cheek 2022). Thus, we modelled the transition from floater to breeding classes as density-dependent. We also assumed breeder carrying capacity was constant across all years which was modelled using a Poisson distribution. Each year floaters had a chance to fill in open spaces created by deceased breeders. Because yearlings are only rarely able to fill such spots (Desrosiers et al. 2021), we first used a binomial distribution to model territory acquisition by surviving floaters from the previous year (i.e. the breeder carrying capacity minus the number of surviving breeders). We used an informative prior distribution (mean = 0.83; beta: α = 10, β = 2) for the probability of any post-yearling floaters acquiring an available territory based on previous work (Bakker et al. 2020). This meant most open territories were taken by older floaters and we assumed that surviving yearlings floaters would fill in any remaining open territories. We also compared our results to results from a density independent IPM to evaluate whether our assumptions about density dependent territory acquisition biased our conclusions (Appendix S1).

### Model Implementation

We ran the population model with 3 Markov chain Monte Carlo (MCMC) chains for 500,000 iterations using default samplers in the package *nimble* in program R (de Valpine et al. 2017; NIMBLE Development Team 2021; R Core Team 2021), discarding the first 50,000 samples and thinning the remaining samples by a factor of 20. We assessed MCMC convergence using Gelman-Rubin diagnostics (*R̂* < 1.1). We ensured models fit adequately by using posterior predictive checks for each dataset (Appendix S2). We also compared our population size estimates with independent indices of abundance from annual Christmas Bird Counts using Pearson’s correlation coefficient.

### A Posteriori Analyses

#### Demographic drivers of population growth

We assessed the effect each vital rate, including population density, has on population growth using Pearson’s correlation coefficient. Because population growth rate is derived from our estimates of population density, we further tested the importance of density dependence at the population-level against a null reference distribution (Schaub & Kéry 2022). We also calculated the variance in each vital rate across years. We propagated all the uncertainty in each parameter by calculating this value across the entire posterior distribution.

#### Ecological drivers of vital rates and population growth

We examined the effect of precipitation, acorn abundance, and population density on vital rates across years *a posteriori* using multiple linear regression (Fig. 2). This approach greatly simplified our IPM structure and allows us to fully propagate uncertainty from the IPM to subsequent analyses. All models included coefficients for population density, acorn abundance, and precipitation in the current and previous years. In survival models, we also tested for an effect of summer maximum and winter minimum temperature since extreme temperatures can increase metabolic costs and potentially impact survival. In the fecundity model, we also included several additional tests. First, we included fixed effects for acorn abundance in the following and previous fall to test two a priori hypotheses. Acorn abundance in the previous fall may affect fecundity through carry-over effect from body condition or caching success. Acorn abundance in the subsequent fall may be perceived by parents because acorn production depends on lagged precipitation and temperatures that are also directly experienced by jays (Koenig et al. 2016). Thus, jays may increase reproductive effort in years when juveniles will have abundant food in their first fall. We also included the density of island foxes, a key nest predator, as a fixed effect. Finally, we ran a model for population growth and included all ecological factors with strong associations (i.e. the proportion of the posterior that is the same sign as the estimate; pd > 0.9) with at least one vital rate. We compared all possible combinations of covariates using AICc and retained model averaged estimates for each variable. We propagated uncertainty in each parameter by calculating these estimates across the entire posterior distribution (i.e. for every iteration of the MCMC chain).

**Fig. 2:**
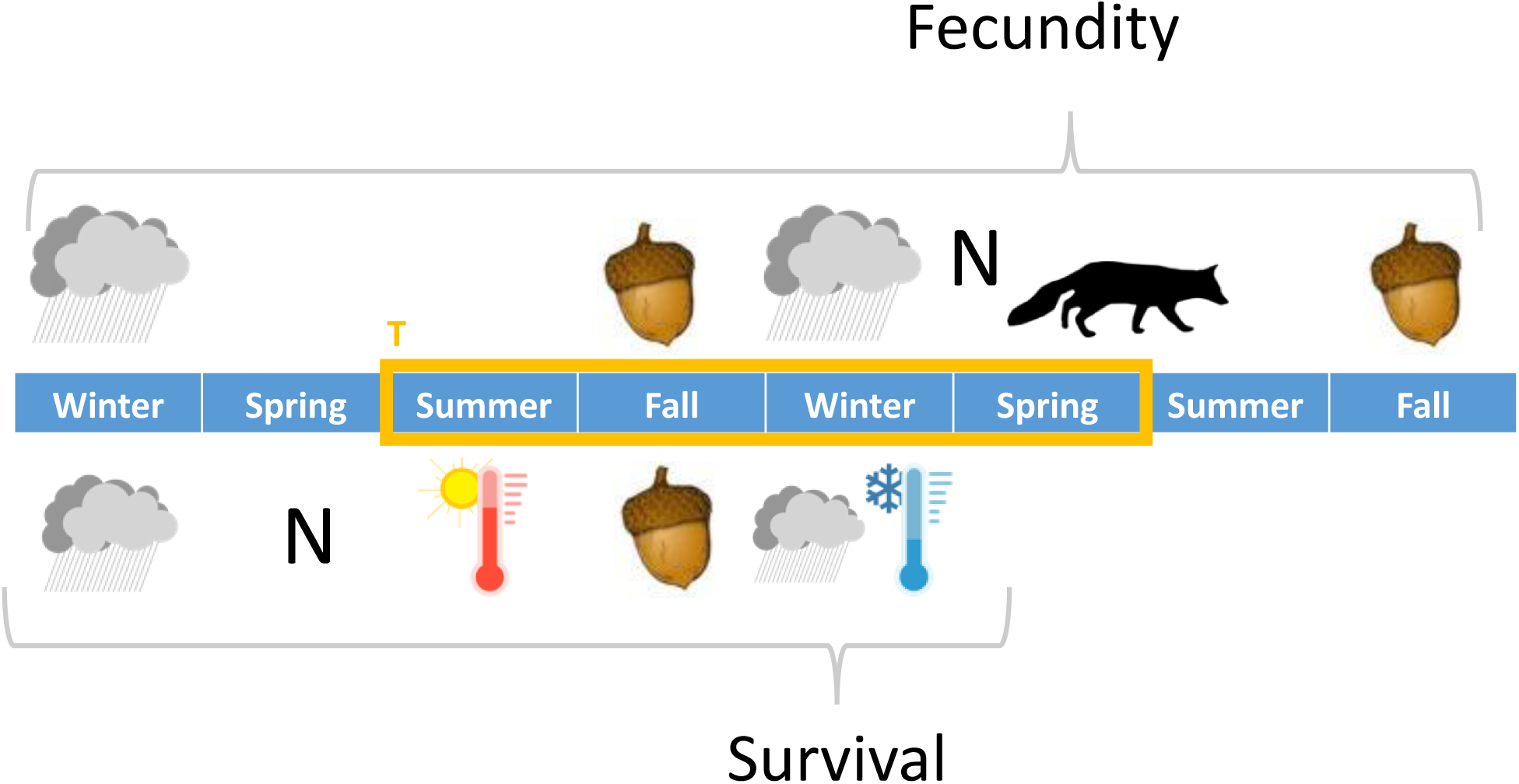
Diagram showing the relative timing of ecological factors compared to survival and fecundity parameters for each bio-year (T). See main text for details.

## RESULTS

We found evidence that island-wide population size of Island Scrub-Jays showed little inter-annual variability, and likely increased between 2009 and 2019 (Fig. 3A; median percentage change in population size = 23.6%; 95% HDI = -23.6 – 86.2%; median number = 378; 95% HDI = -382 – 1136; pd = 0.849). Breeders had relatively high and consistent annual survival rates across years (Fig. 3B; mean = 0.89, sd = 0.06, range of annual means = 0.75 - 0.96) compared with non-breeding floaters (Fig. 3B; mean = 0.83, sd = 0.10, range of annual means = 0.63 - 0.96). The expected number of fledglings produced per territory varied substantially among years but was consistently less than 1 (Fig. 3C; mean = 0.31, sd = 0.14, range of annual means = 0.11 - 0.55). Accordingly, annual population growth rate was weakly associated with temporal variation in breeder survival (Fig. 4A; median = 0.39; 95% HDI = -0.12 – 0.80) but showed relatively strong associations with floater survival (Fig. 4B; median = 0.70; 95% HDI = 0.36 – 0.93) and fecundity (Fig. 4C; median = 0.67; 95% HDI = 0.19 – 0.93).

**Figure 3:**
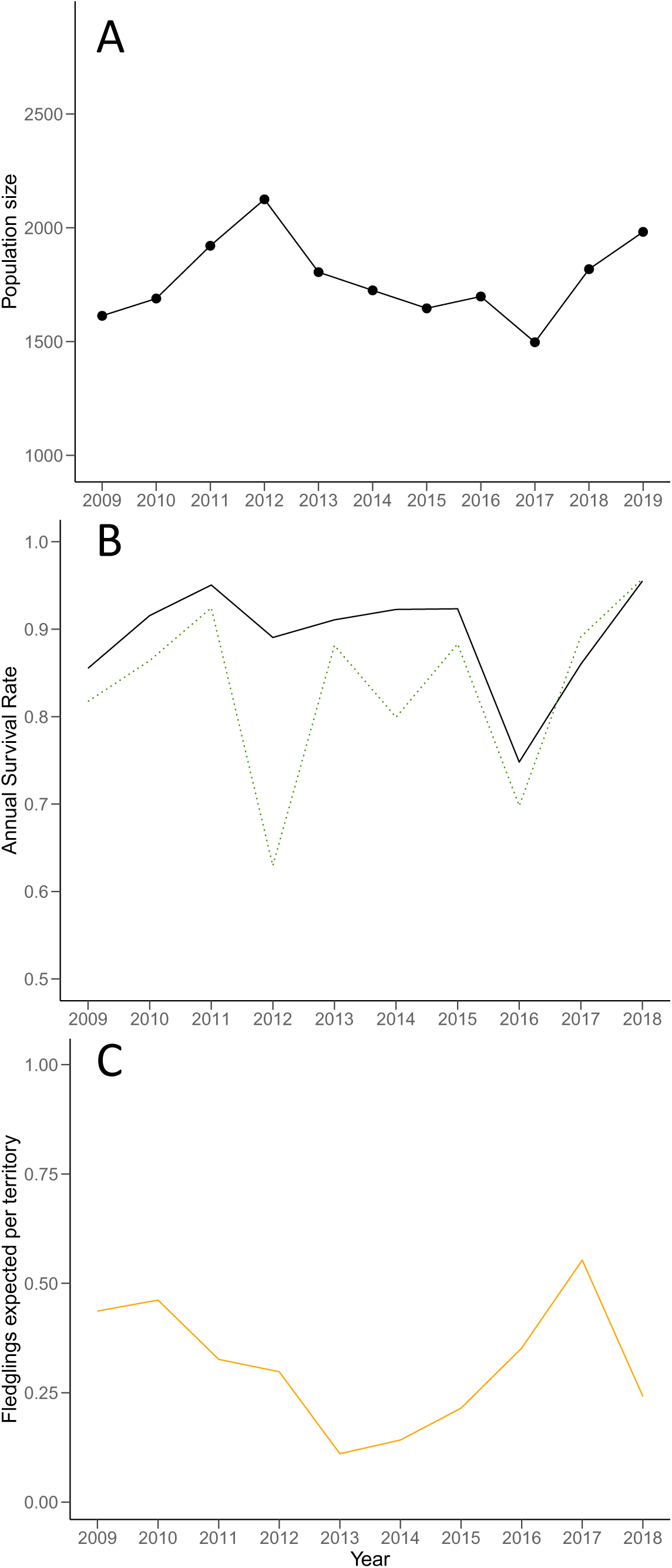
**A)** Island-wide population size, **B)** breeder (grey, solid line) and non-breeder (green, dotted line) annual survival, and **C)** fecundity for Island Scrub-Jays from 2009-2019. Lines depict posterior medians and shaded areas depict 95% highest posterior density intervals.

**Figure 4:**
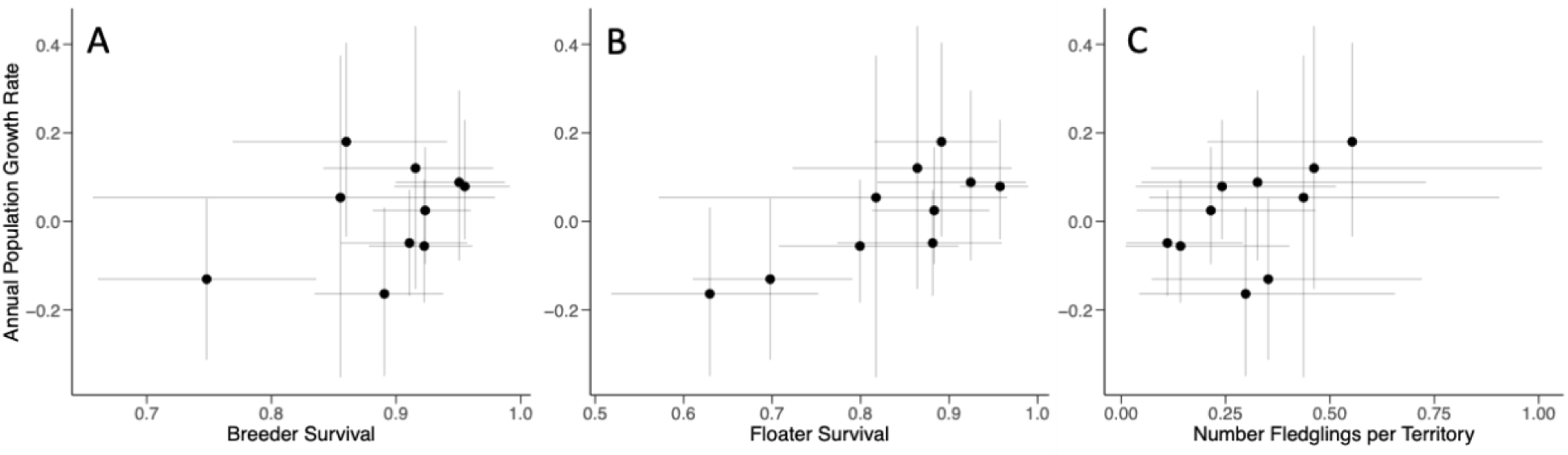
Relationships between annual population growth rate and **A)** breeder survival, **B)** floater survival, and **C)** fecundity for Island Scrub-Jays from 2009-2019. Points depict posterior medians and errorbars depict 95% highest density intervals.

Island-wide population size did not strongly affect breeder survival (Fig. 5A; median = 0.13; 95% HDI = -0.39 – 0.59; pd = 0.69). However, we did find support for negative density dependence acting on floater survival (Fig. 5B; median = -0.34; 95% HDI = -0.71 – 0.14; pd = 0.92), and fecundity (Fig. 5C; median = -0.39; 95% HDI = -0.79 – 0.09; pd = 0.93). Population growth rate was also negatively correlated with island-wide population size across years (Fig. 5D; median = -0.58; 95% HDI = -0.85 – -0.34; pd > 0.99); support for this inference remained relatively high after comparison with a null reference distribution (pd = 0.80). Population size fluctuated around a median carrying capacity of 1823 breeders over the course of the study (95% HDI = 1384 – 2332). We found similar patterns and support for density dependence from our density independent IPM (Appendix S1) and in models accounting for environmental factors (Table 1).

**Figure 5:**
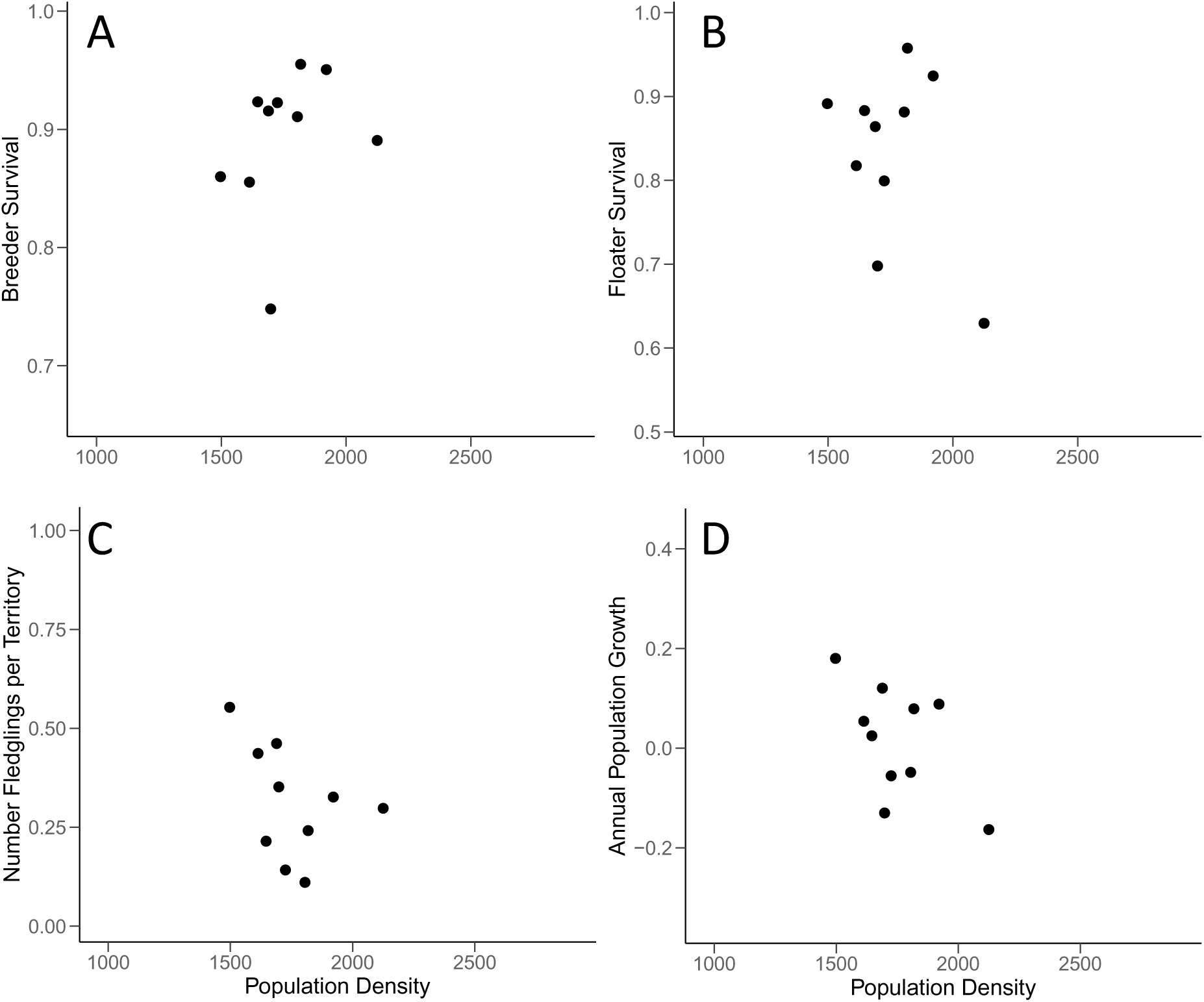
**A**) Relationships between island-wide population size and **A)** breeder survival, **B)** non-breeder survival, **C)** fecundity, and **D)** annual population growth for Island Scrub-Jays from 2009-2019. Points depict posterior medians and errorbars depict 95% highest density intervals.

**Table 1:** Model averaged regression coefficients across entire posterior distribution for each demographic parameter. We show median and 95% highest density intervals for model averaged coefficients for each ecological factor. The probability that a posterior distribution is strictly negative or positive pd> 0.95, pd > 0.90, and pd > 0.85, are depicted in bold with **, *, and **^+^** respectively. We include results from both breeder survival models based on all years and without 2016 (see text for details).

| Factor | Population Growth Rate | Breeder Survival | Breeder Survival (w/o 2016) | Floater Survival | Fecundity |
| --- | --- | --- | --- | --- | --- |
| Population Density | **-0.18 (-0.88 – 0.01) \*\*** | 0.005 (-0.25 – 0.25) | 0.005 (-0.20 – 0.27) | **-0.03 (-0.37 – 0.09) .** | **-0.04 (-0.58 – 0.07) +** |
| Acorn Abundance | -0.004 (-0.13 – 0.12) | **-0.04 (-0.27 – 0.02) \*** | 0.003 (-0.10 – 0.15) | **0.01 (-0.01 – 0.06) .** | 0.007 (-0.09 – 0.27) |
| Previous Year Acorn Abundance | **0.18 (-0.02 – 0.84) \*\*** | - | - | - | **0.07 (-0.04 – 0.72) \*** |
| Precipitation | **0.06 (-0.03 – 0.57) \*** | 0.003 (-0.05 – 0.08) | -0.007 (-0.16 – 0.08) | **0.02 (-0.02 – 0.12) \*** | **0.06 (-0.03 – 0.79) \*** |
| Previous Year Precipitation | - | 0.02 (-0.03 – 0.16) | 0.08 (-0.07 – 0.15) | 0.007 (-0.04 – 0.07) | 0.004 (-0.11 – 0.30) |
| Winter Min. Temperature | -0.009 (-0.53 – 0.14) | **-0.04 (-0.27 – 0.02) \*\*** | -0.007 (-0.12 – 0.06) | 0.005 (-0.05 – 0.07) | - |
| Summer Max. Temperature | - | 0.004 (-0.06 – 0.09) | -0.02 (-0.37 – 0.06) | 0.01 (-0.05 – 0.14) | - |
| Island Fox Density | - | - | - | - | -0.01 (-0.30 – 0.09) |

Precipitation, temperature, acorn abundance, and the density of island foxes differed among years, but the effects on vital rates varied (Table 1). Annual precipitation was positively associated with fecundity (pd = 0.92) and floater (pd = 0.87) survival, but not breeder survival (pd = 0.59). Fecundity was also positively associated with the abundance of acorns the previous (pd = 0.92), but not the subsequent (pd = 67), fall. Breeder survival was negatively associated with acorn abundance (pd = 0.94), minimum winter temperature (pd = 0.96), and showed a weak positive association with precipitation in the previous year (pd = 0.84). However, these patterns were driven by a single outlier year of low breeder survival (2016). When this year was removed from the data, no strong relationship existed between breeder survival and acorn abundance (median = 0.003; 95% HDI = -0.10 – 0.15; pd = 0.57) or minimum winter temperature (median = -0.007; 95% HDI = -0.12 – 0.06; pd = 0.69), and the only potentially meaningful pattern was a weak negative relationship with maximum summer temperature (median = -0.02; 95% HDI = - 0.38 – 0.06; pd = 0.80). Temporal variation in population growth was negatively related to population density (pd = 0.98) and positively associated with precipitation (pd = 0.95) and acorn abundance the previous fall (pd = 0.96).

## DISCUSSION

Climate change and other anthropogenic impacts are expected to have major implications for population dynamics, but the degree to which current conditions are surpassing demographic buffering mechanisms is unclear. Here we provide evidence that Island Scrub-Jay population size has increased slightly over the last decade and appears to be relatively stable despite record high temperatures and a significant multiyear drought. Additionally, we found evidence for both density-dependent population regulation acting primarily through early life stages and an important role for environmental effects on key vital rates. Overall, our results suggest jays have been capable of buffering recent environment variation and provide useful information about the ecological drivers of population dynamics in island species.

A key aspect of a population’s ability to persist under changing environments is its ability to self-regulate. We found evidence for density dependent population regulation among Island Scrub-Jays at the population-level that seemed to act primarily via effects on fecundity and floater survival (Fig. 5; Table 1). This form of density dependence is common among species with slower life history strategies and longer lifespans and likely helps buffer more important vital rates like breeder survival from environmental variation (Eberhardt 2002, Galillard et al 1998). Though we did not find evidence that island fox density had any impact on fecundity through nest predation, it seems likely that conspecific nest predation by jays may decrease nest and juvenile survival in high density years. Jays have even been observed consuming their own eggs after perceiving a nest predator finding their nest which may also drive density dependence in fecundity (Caldwell et al 2012). Regardless of the specific mechanism, density dependence in fecundity and floater survival seem to be important for population regulation in this species.

Our models also assumed that recruitment of floaters into the breeding population would be density-dependent. We decided to integrate this assumption into the model based on our observation of territory stability, territory acquisition by floaters, and behavioral interactions between breeders and floaters in the field. This assumption did not bias our conclusions about density dependence in vital rates or at the population level because our density-independent model did not make this assumption and produced nearly identical results (Appendix S1). Nonetheless, more detailed study of this process and the biology of non-breeding floaters in general would be useful in testing this idea more explicitly. For example, territory acquisition may be related to natal-habitat preferences and individual variation in traits like body size (Desrosiers et al. 2021; Cheek et al. 2024). Multistate-survival models may be able to provide estimates of territory acquisition from our data and collecting data on the movements, behaviors, and abundance of the non-breeding floater subpopulation may help us understand this process in more detail. Directly monitoring floater subpopulations may also prove useful for more rapidly identifying declines among floaters which may provide warning about impending declines among territorial breeders and in the population more broadly (Porter & Coulson 1987; Klomp & Furness 1992; Hunt 1998).

Precipitation had major effects on population dynamics in Island Scrub-Jays. We found positive effects of precipitation on fecundity and floater survival, but negligible effects on breeder survival (Table 1). Precipitation-driven variation in these vital rates yield increased population growth rates. Precipitation is also intimately linked with breeding behavior and fecundity in many species in other semi-arid ecosystems (Gibbs & Grant 1987; Morrison & Bolder 2002; Sofaer et al. 2014). For example, breeding failure in Orange-crowned Warblers, which occur on Santa Cruz Island and several other nearby islands, is greatly reduced in years with more rainfall (Sofaer et al 2014). In addition to short-term variation in rainfall, persistent drought may also have critical implications for population growth over longer time-scales. In our system, wet winters can have large impacts on food supply during the spring, and also influence acorn production the following summer (Koenig et al. 1996). Acorns provide food for jays throughout subsequent fall and winter and are positively associated with fecundity the following year (Table 1). Rainfall and especially persistent drought can also impact affect the distribution and quality of habitat (Taylor et al. 2020) with implications for the population’s carrying capacity.

Despite breeder survival being relatively invariant across years, reduced breeder survival in 2016 was associated with high acorn abundance and warmer winter temperatures (Fig. 3b; Table 1). Neither acorn abundance nor winter temperature scaled up to influence population growth and these relationships were driven primarily by a single year. Nonetheless, the acorn results were surprising because we expected greater availability of a key food resource and milder winters to create favorable conditions for survival. This decline in breeder survival also coincided with the last year of a significant multiyear drought and may also be explained by cumulative effects of drought on breeder survival. Delayed demographic responses to drought have been observed among passerines in similar chaparral habitat on the adjacent mainland and, in general, species at higher trophic levels may have the most lagged response times (Sakai 2016; Prugh et al. 2018). Breeding Bird Survey (BBS) data from the mainland suggest that California Scrub-Jay (*Aphelocoma californica*) abundance declined sharply in 2016 (∼9%; Sauer et al. 2021) hinting that declines in breeder survival in Island Scrub-Jays could have been caused by the persistent drought or other regional factors. Given the relative stability of breeder survival, confidently distinguishing among different ecological drivers would require continuous study of breeder survival over even longer time scales. In any case, since the relationships between breeder survival, winter temperature, and acorn abundance were driven by a single year with low breeder survival and did not influence population growth (Table 1), it should perhaps be interpreted with caution.

We found more compelling evidence that acorn abundance influences fecundity in the following spring, but no evidence that jays plastically adjusted reproductive output to match expected acorn crops the following fall. Such predicted adaptive responses are theoretically possible because oaks initiate buds that ultimately yield acorns as long as 18 months prior to their harvest by jays (*unpublished data*; Fleurot et al. 2023). Other seed-eating species, such as American (*Tamaiasciurus hudsonicus*) and European (*Sciurus vulgaris*) red squirrels, increase reproductive output prior to seed masting events in conifers (Boutin et al. 2006). It may be that energetic resources during the spring as nestlings and fledglings may be more important for the growth and recruitment of individual young (Desrosiers et al. 2021). Increased consumption and caching of acorns in the fall could both lead to greater fecundity the next breeding season. Greater consumption of acorns in the time leading up the breeding season could yield improved body condition and fuel egg production, parental care, and ultimately reproductive success. Greater consumption of cached acorns during breeding might also help offset energetic cost of parental care and facilitate successful breeding (Koenig & Mumme 1987). In any case, these patterns suggest that ecological conditions experienced during the non-breeding season can carry-over to impact reproductive success during the follow breeding season and highlight the importance of full-annual cycle research in even the least migratory species.

We found evidence for a relatively stable and even increasing population from 2009-2019 and have seen consistent territory occupancy across all our breeding plots. Despite risks that jays continue to face; this should come as welcome news to managers and suggests a relatively resilient populations under current environmental conditions. Recent work from a field study of jays in the center of the island suggested that jays had suffered alarming declines in breeder survival and territory acquisition by floaters from 1999-2006 (Mosher et al. 2021). These declining vital rates suggest severe decreases in breeding density of the study population during those years but did not seem to be associated with variation in winter precipitation, the eradication of feral sheep or pigs, or WNV (Mosher et al. 2021). A rapidly expanding invasive wild turkey population coincided with this decline and may have impacted jay floater survival through competition for acorns and other food (Morrison et al. 2016; Mosher et al. 2021). Turkeys were concentrated in two areas on the island including near the focal population for this study. In contrast, our work took place at a number of sites across the island and followed the eradication of turkeys (Morrison et al. 2016). The patterns we found suggest that the declines in vital rates and population size from 1999-2006 have either stopped or were limited to specific location studied during that time.

### Implications for Management

Despite evidence for relative population stability during this study, Island Scrub-Jays remain vulnerable to extinction in the long-term due to their limited range and small population size (Morrison et al. 2011; Bakker et al. 2020). Climate change is expected to increase temperatures and exacerbate drought in southern California in the future (Cayan et al. 2008).

More frequent and severe droughts may reduce the number of non-breeding floaters available to take over breeding territories in the long-term and may also impact breeder survival directly under especially prolonged or extreme conditions. Warming temperatures and reduced rainfall due to climate change will also likely exacerbate the risk of heat-related mortality, wildfire or novel infectious disease on the island. For example, a severe wildfire on the island could quickly destroy the majority of breeding habitat for the species (Morrison et al. 2011). Warmer temperatures may also facilitate the establishment and spread of diseases like West Nile virus which is endemic on the mainland a mere 32 km away (Boyce et al. 2011). Island Scrub-Jays are thought to be especially sensitive to WNV given their relative exposure to other pathogens and the dramatic population declines experienced by related species on the mainland (Koenig et al. 2007; Morrison et al. 2011). Either fire or WNV could rapidly cause catastrophic decline in vital rates and population size of Island Scrub-Jays and pose a true existential threat. Overall, the range of long-term and potentially more immediate risks faced by Island Scrub-Jays require careful monitoring and thoughtful management.

Given the relatively high severity and likelihood of these threats, proposed, proactive conservation actions for island scrub-jays include management of a captive population, maintaining a sub-population on Santa Cruz Island vaccinated against WNV, and re-establishing a second wild population of jays on Santa Rosa Island (Morrison et al. 2011). Previous work has suggested that in both vaccination and reintroduction to Santa Rosa would likely reduce global extinction risk substantially (Bakker et al. 2020). The fact that we found evidence for density dependence and breeding population saturation suggests that removal of wild jays, especially floaters, from Santa Cruz Island to establish these secondary populations would have little effect on overall population size. Reintroduction of jays to Santa Rosa Island could also aid in the restoration of oak and pine habitat there because of the long-distance seed dispersal services jays provide (Pesendorfer et al. 2018). More research would help improve the likelihood of a successful reintroduction to Santa Rosa. For example, studying habitat selection and demography of jays living in the wind-blown chaparral on the far western portion of Santa Cruz Island that is more similar in habitat and climate to Santa Rosa would help understand population dynamics after reintroduction. Additional study of jay diet could also help allay concerns about predation of vertebrates by jays on Santa Rosa. Undoubtedly, some uncertainty will remain about the outcome of a reintroduction. However, proactive approaches can greatly reduce the risk of fast-moving threats like WNV to imperil Island-Scrub Jays.

Understanding the demographic mechanism underlying population dynamics is important for assessing contemporary and future effects of climate change. Our findings are consistent with theory and recent empirical work suggesting long-lived species should have population dynamics driven most strongly by fecundity and non-breeder survival (Pfister 1998; Hilde et al. 2020). We were also able to identify environmental variables most likely to impact populations under normal and more novel conditions in the coming decades. Coupling long-term fieldwork with integrated population modelling to gain insight on demographic and environmental influences on population dynamics promises to advance population ecology and conservation management into an uncertain future.

## ACKNOWLEDGEMENTS

We thank anonymous reviewers for helpful comments. We are also grateful for support from The Nature Conservancy, Channel Islands National Park, and the U. S. National Science Foundation (DEB-1754821). We would especially like to thank Annie Littlle, Dylan Moe, Jen Baker, Amy Parks, Andrew Yamagiwa, Scott Meyler, Hannah Horowitz, and Sarah Hays. This work was conducted under the auspices of NZP ACUC protocol SI-22023 (Ecology of landbirds on the California Channel Islands). Any use of trade, product, or firm names is for descriptive purposes only and does not imply endorsement by the U.S. Government.

## AUTHOR CONTRIBUTIONS

JCM, TSS conceived the idea for the study. TSS, MP, and designed and refined the field methodology. TSS, MP, and JCM collected the data. JCM created the modelling framework. JCM led the writing of the manuscript. All authors contributed critically to the drafts and gave final approval for publication.

## Appendix S1 Density-independent IPM

Our main IPM explicitly assumes that non-breeding floaters acquire territories in a density dependent manner. This assumption is based on observations in the field and previous work in the system (Desrosiers et al. 2021; Collins & Corey 1994). Nonetheless, we directly tested these assumptions comparing results from a complementary density-independent IPM. This density-independent IPM used the same sub-model structure for population count, survival, and fecundity data, but had a slightly different population model at its core. We repeated all *a posteriori* analyses using the methods detailed in the main text.

### Methods

#### Population model

As in our main IPM, we modeled island-wide population size in 2009 using a negative binomial prior distribution parameterized to match estimates from previous work (1705; 1212-2369 95%CI; Sillett et al. 2012). We modelled the number of surviving juvenile (from fledge until the subsequent breeding season), floater, and breeder survival using binomial distributions and survival parameters. Territory acquisition was modelled using binomial distributions. We modelled the probability of acquiring a territory for first-year and after-first-year floaters using beta prior distributions. These distributions were parameterized using previous work in the system. This model structure does not limit the number of breeding birds on the island and allows floaters to continuously acquire territories regardless of the current number of breeding birds already present.

#### Model Implementation

We ran the population model with 3 MCMC chains for 500,000 iterations using default samplers in the R package *nimble* (NIMBLE Development Team 2021). We discarded the first 50,000 samples and thinned the remaining samples by a factor of 20. We assessed MCMC convergence using Gelman-Rubin diagnostics (*R̂* < 1.1). We ensured models fit adequately using posterior predictive checks for each dataset (see Appendix S2 for details). We again compared population size estimates with independent data from annual Christmas Bird Counts using Pearson’s correlation coefficient. These CBC data were which were collected several months prior to the breeding season in all years except 2010, 2014, and 2018.

### Results

The density-independent model predicted similar patterns and increases in population size from 2009 to 2019 as our main IPM (Fig. S1; median percentage = 28.5%; 95% HDI = -20.1 – 86.3%; median number = 430; 95% HDI = -319 – 1308; pd = 0.876). Temporal patterns in population growth rate (Pearson’s r: median = 0.53; 95% HDI = -0.17 – 0.88; pd = 0.942), breeder survival (Pearson’s r: median = 0.67; 95% HDI = 0.01 – 0.96; pd = 0.976), floater survival (Pearson’s r: median = 0.68; 95% HDI = 0.12 – 0.94; pd = 0.993), and fecundity (Pearson’s r: median = 0.44; 95% HDI = -0.14 – 0.84; pd = 0.932) from the density-independent model were are positively correlated with patterns from our main IPM.

The correlation between each vital rate and population growth (Fig. S2) and population density (Fig. S3) was similar between our main IPM and the density-independent model. Estimates of island-wide carrying capacity from each model were also broadly overlapping (density-independent model K: median = 1741; 95% HDI = 1045 – 2762; main IPM K: median = 1822; 95% HDI = 1384 – 2332). Support for density dependence at the population remained about as strong in the density-independent model (pd = 0.82) as our main IPM (pd = 0.80) after comparison with a null reference distribution.

We found similar patterns in the relationship between population vital rates and environmental variables (i.e. precipitation, acorn abundance, temperature, and island fox density) and between the density-independent model and our main IPM (Table S1) suggesting that our assumptions about density dependent territory acquisition do not bias our conclusions.

**Figure S1:**
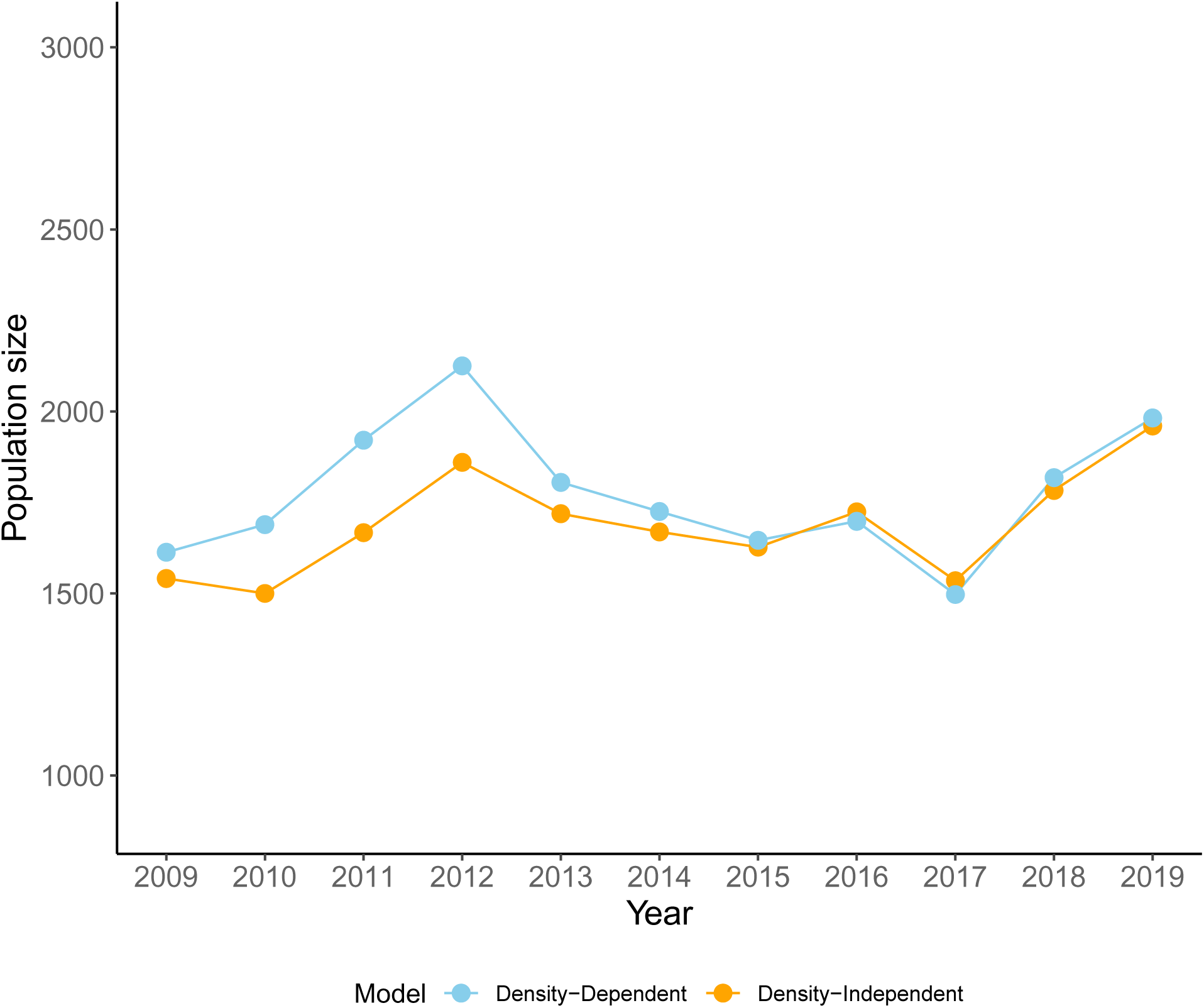
Island-wide population size for Island Scrub-Jays from 2009-2019 from our main IPM with density dependent territory acquisition (blue) and the density-independent model (orange). Lines depict posterior medians and shaded areas depict 95% higher density intervals.

**Figure S2:**
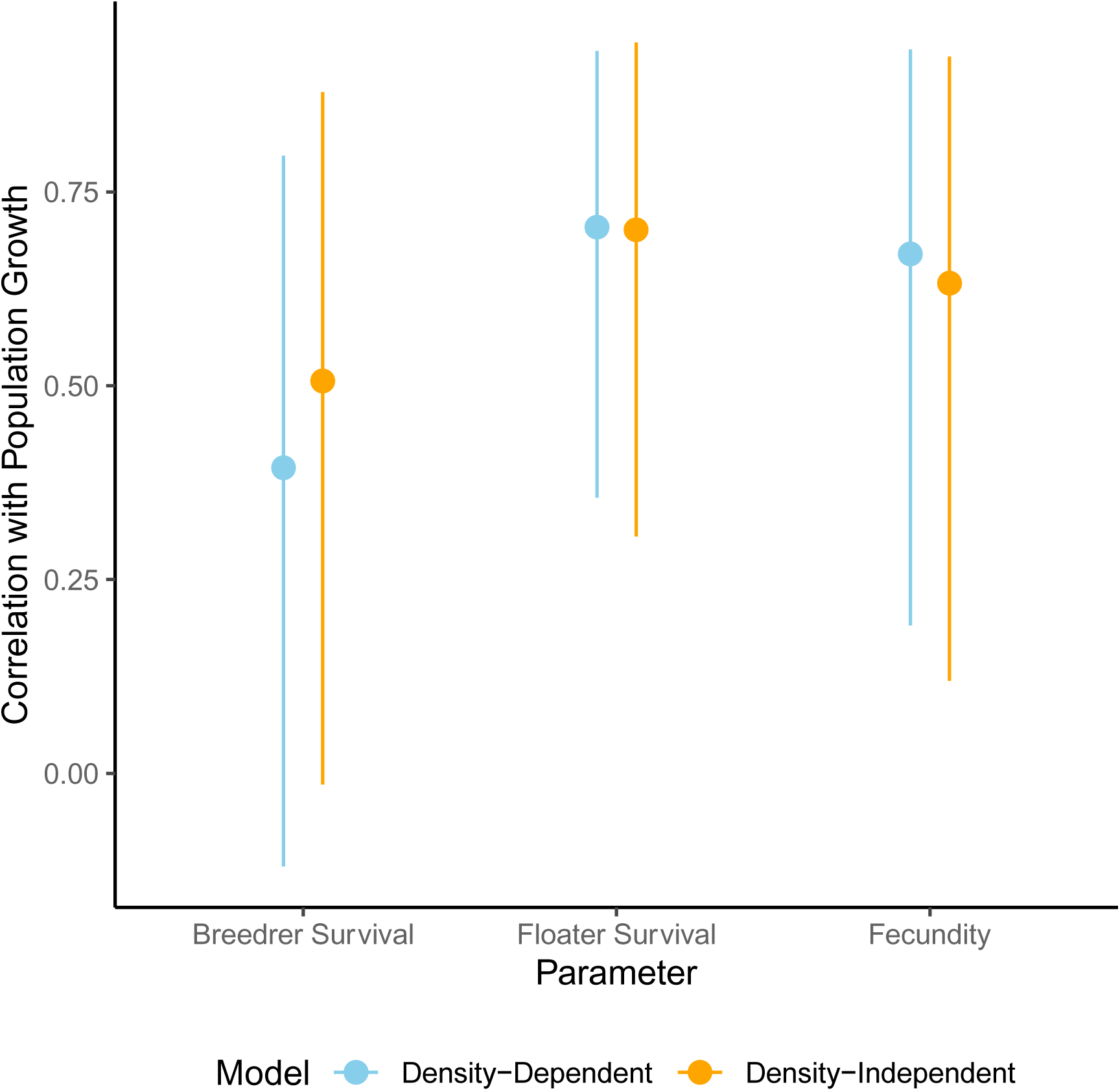
The relationship between population vital rates and population growth for our main IPM with density dependent territory acquisition (blue) and the density-independent model (orange). Points depict median and 95% highest density intervals.

**Figure S3:**
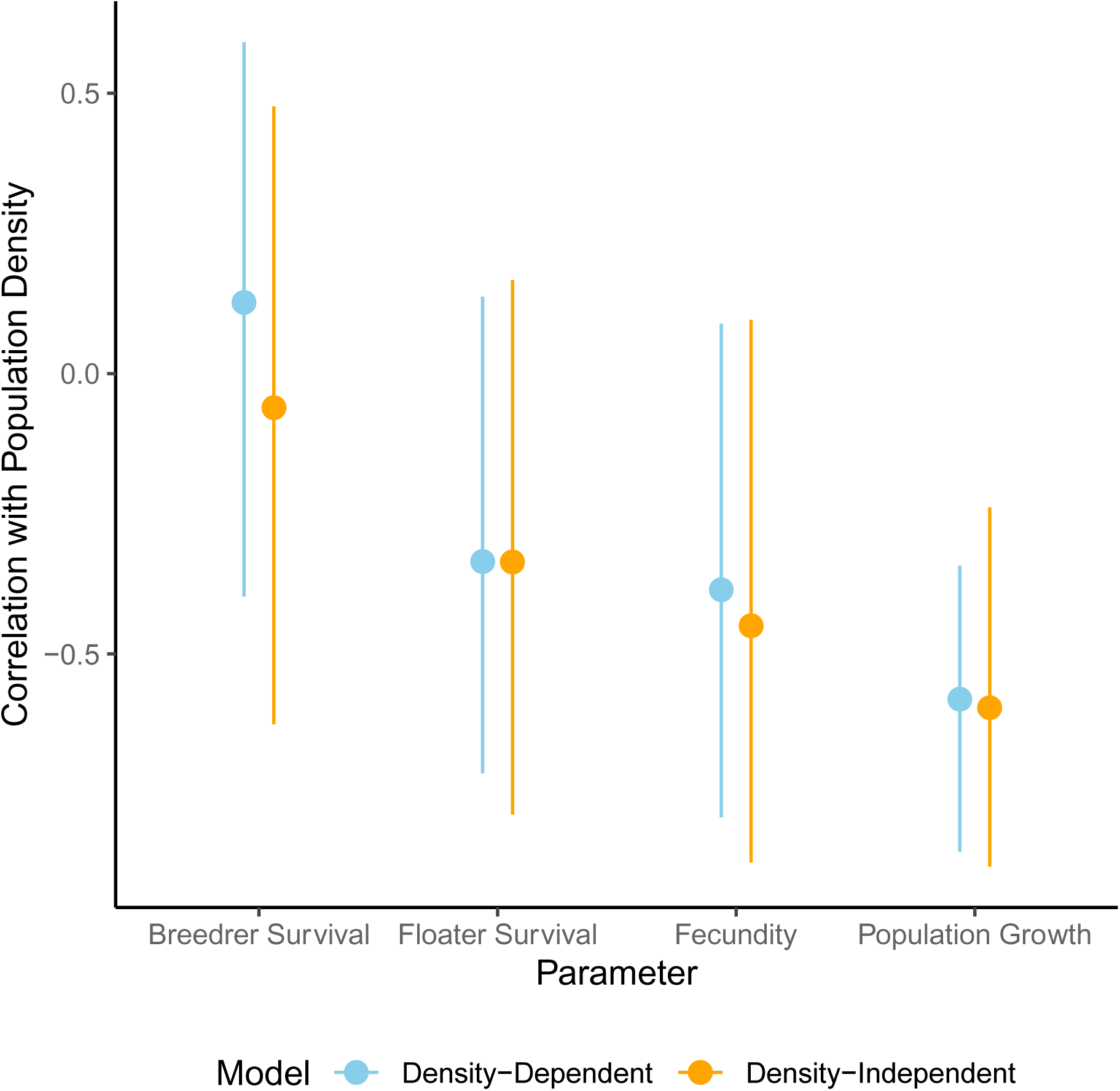
The relationship between population vital rates and island-wide population density for our main IPM with density dependent territory acquisition (blue) and the density-independent model (orange). Points depict median and 95% highest density intervals.

**Table S1:** Model averaged regression coefficients across entire posterior distribution for each demographic parameter. We show median and 95% highest density intervals for model averaged coefficients for each environmental factor. Estimates with pd > 0.95, pd > 0.90, and pd > 0.85 are depicted in bold with **, *, **^+^** respectively.

| Environmental Factor | Population Growth Rate | Breeder Survival | Floater Survival | Fecundity |
| --- | --- | --- | --- | --- |
| Population Density | -0.90 (-5.48 – 3.93) | -0.004 (-0.43– 0.21) | -0.02 (-0.47 – 0.10) | **-0.06 (-0.78 – 0.07)** <sup>+</sup> |
| Acorn Abundance | -0.07 (-2.21 – 2.06) | **-0.04 (-0.276– 0.02)** * | **0.01 (-0.01 – 0.06)** . | 0.005 (-0.11 – 0.26) |
| Previous Year Acorn Abundance | 0.09 (-4.35 – 4.21) | - | - | **0.05 (-0.05 – 0.70)** <sup>+</sup> |
| Precipitation | 0.47 (-2.24 – 2.74) | 0.003 (-0.05 – 0.08) | **0.02 (-0.02 – 0.11)** <sup>+</sup> | **0.06 (-0.04 – 0.79)** * |
| Previous Year Precipitation | 0.40 (-2.87 – 3.51) | **0.02 (-0.03 – 0.18)** <sup>+</sup> | 0.007 (-0.04 – 0.07) | 0.009 (-0.10 – 0.38) |
| Winter Min. Temperature | -0.10 (-4.09 – 4.18) | **-0.04 (-0.26 – 0.02)** ** | 0.006 (-0.03 – 0.05) | - |
| Summer Max. Temperature | 0.23 (-1.54 – 1.82) | 0.004 (-0.05 – 0.08) | 0.01 (-0.04 – 0.14) | - |
| Island Fox Density | 0.14 (-2.23 – 2.49) | - | - | -0.02 (-0.34 – 0.10) |

## Appendix S2 Model checking

### Posterior Predictive Checks

We ensured models fit adequately using posterior predictive checks for each dataset. Posterior predictive checks compare some discrepancy or summary values of the real dataset with the same value from a dataset simulated from each MCMC iteration of the model. We then calculated the percentage of summary statistics were less than the summary statistic based on the real data. This yielded a Bayesian P-value where P>0.9 or P<0.1 indicates poor model fit.

For the abundance sub-model, we used the mean absolute percentage error (MAPE; Besbeas & Morgan 2014). We also compared the overall mean of counts across years. Both summary statistics suggested that the abundance sub-model adequately fit the data (P_MAPE_ = 0.60; P_Mean_ = 0.46).

For the fecundity sub-model, we used the Freeman-Tukey statistic (Brooks et al. 2000). We also compared the overall mean number of fledglings across years and the number of territories producing no fledglings. The fecundity sub-model fit the data adequately based on each discrepancy measures (P_Freeman-Tukey_ = 0.27; P_Mean_= 0.33; P_Zeros_= 0.54).

For the survival sub-model, we calculated the Goodness-of-fit using the combined test for Cormack-Jolly-Seber models (i.e. Test 1; Giminez et al. 2017). We calculated this value for each group (sex, breeding status, first-year bird) and found the mean to determine the overall fit. The survival sub-model fit the data adequately (P = 0.14).

### Comparison with independent dataset

We also compared island-wide population size estimates with independent data from annual Christmas Bird Counts using Pearson’s correlation coefficient. These CBC data were collected several months prior to the breeding season in all years except 2010, 2014, and 2018 when no CBC was conducted on the island (National Audubon Society 2022). We calculated the correlation for each MCMC iteration. Overall, despite the noise and slightly altered timing of the CBC data, the model’s estimates of island-wide population size were weakly associated with the CBC abundance indices across years (r = 0.19; 95% HDI = -0.32 – 0.76; pd = 0.74).

